# Peripheral tolerance to insulin is encoded by mimicry in the microbiome

**DOI:** 10.1101/2019.12.18.881433

**Authors:** Arcadio Rubio García, Athina Paterou, Mercede Lee, Hubert Sławiński, Linda S. Wicker, John A. Todd, Marcin Ł. Pękalski

## Abstract

How organisms achieve sustained peripheral tolerance throughout their lifetime, a correct immune discrimination between self and non-self, remains poorly understood. Host-microbiome interactions carry fundamental information that facilitates this process. We hypothesize that commensal microbes are under evolutionary pressure to develop epitopes that, when presented along with other antigens from their own bacterial community, lead to an overall tolerogenic self classification by the host immune system. Hosts, which have co-evolved with commensals, may rely on mimotopes, bacterial epitopes that are indistinguishable from key self epitopes, as a homeostatic feedback mechanism to establish and maintain tolerance. Using a probabilistic sequence model of peptide mimicry, we show that the gut microbiome contains a set of genes that are likely to trigger identical immune responses to insulin B 9–25, a widely distributed self epitope across tissues and the primary autoantigen in type 1 diabetes. Similarities in the antigen receptor sequences determined from CD4 T cells reacting to insulin epitopes and mimotopes provide experimental evidence for mimicry. All predicted high posterior probability mimotopes belong to the transketolase superfamily, an enzyme that allows efficient harvest of commensal-derived sugar polymers and dietary fibre, an advantage during host colonisation. Microbial transketolase upregulation during infant weaning coincides in time with the peak in autoantibody development against insulin. Abundance changes in bacterial genera that carry these mimotopes have also been observed to precede disease diagnosis. Our findings suggest gut dysbiosis followed by immune response to insulin mimotopes as a primary cause of type 1 diabetes, and may contribute towards unraveling similar causal patterns in a wide variety of disorders.

## 1 Introduction

The human microbiome plays a fundamental role in health and disease. Early twin studies showed deviations from a core set of gut microbial genes are associated with different physiological states [1]. This result has been extended to a variety of tissues [2] and disorders [3] such as inflammatory bowel disease [4], multiple sclerosis [5], type 1 diabetes [6], type 2 diabetes [7], Alzheimer’s disease [8], and Parkinson’s disease [9].

Westernized societies exhibit dramatic microbiome differences with respect to isolated hunter-gatherer populations, both in terms of diversity and composition, due to the myriad of modern practices that decrease commensal transmission and survival [10]. These include scarce microbiota-accessible carbohydrates related to low-fibre diets [11] or early-life antibiotic treatments [12], providing some cues of a possible causal chain linking environmental changes and increased prevalence of complex disorders via alterations to the microbiome. Germ-free model organisms and fecal transplants provide further mechanistic evidence by inducing [13] or reversing [14], with some limitations, a variety of disease phenotypes.

Dysregulated metabolic pathways do not seem sufficient to explain the numerous associations between microbiome changes and disease, in particular within autoimmune inflammatory disorders. This has prompted revisiting molecular mimicry [15] in the context of symbiosis and the subsequent discovery of several commensal peptides that may mimic key autoantigens in Sjögren’s disease [16], systemic lupus erythematosus [17], multiple sclerosis [18] and antiphospholipid syndrome [19]. The role of commensal mimicry in human physiology remains unclear as most candidates are limited to a reduced number of microbial genes, some mimotopes originate from bacterial orthologs of host proteins, or belong to species with possible pathogenic potential [20, 21]. Moreover, the lack of a precise definition of molecular mimicry at sequence level hinders further progress, with sequence homology being a poor proxy [22].

Since commensal proteomes do not seem to influence central thymic selection through any primary pathways, mimicry could be a ubiquitous evolved strategy of the microbiome to avoid potentially harmful immune responses against microbes and their metabolites that are essential for the health of the host. Hosts may have co-evolved a mechanism to maintain immune homeostasis by exploiting continuous presentation of mimotopes obtained in the microbiota-mucosal interface to, for example, induce peripheral regulatory T cells or carry out a controlled apoptosis of T effector cells.

We postulate that molecular mimicry occurs when all antigen receptors that can be constructed for a given epitope, or a set of epitopes digested and presented from a particular antigen, bind either the original epitope or the mimotope with equivalent affinity. This may happen if and only if all contact points in the receptor-epitope or receptor-mimotope structure contain identical amino acids or some closely related residue. The remaining positions are more prone to tolerate substitutions with similar or sometimes distant amino acids. In general, insertions or deletions may break mimicry. Using this probabilistic definition of sequence mimicry we uncover a large set of bacteria that are likely to trigger identical immune responses to insulin B 9–25, a widely expressed self epitope and the primary autoantigen in type 1 diabetes (T1D), by adapting a region of the transketolase protein superfamily—an enzyme related to the metabolism of fibre upregulated during infant gut development [23].

## 2 Results

### 2.1 The human gut biome harbors potential insulin B mimotopes

In order to assess the existence of commensal mimotopes, we have defined a generative process that models the mixture of sequences that exhibit some homology, or significant similarity [24], plus those with mimicry potential to a self epitope of choice. The core idea is to sample two different motifs for each class of sequences, homologs and mimotopes, from position-specific prior distributions. Then, for each sequence, to sample its class and each residue from the previously constructed motifs, which now act as distributions, conditioned on the class.

#### Algorithm 1 Hierarchical mixture mimicry model

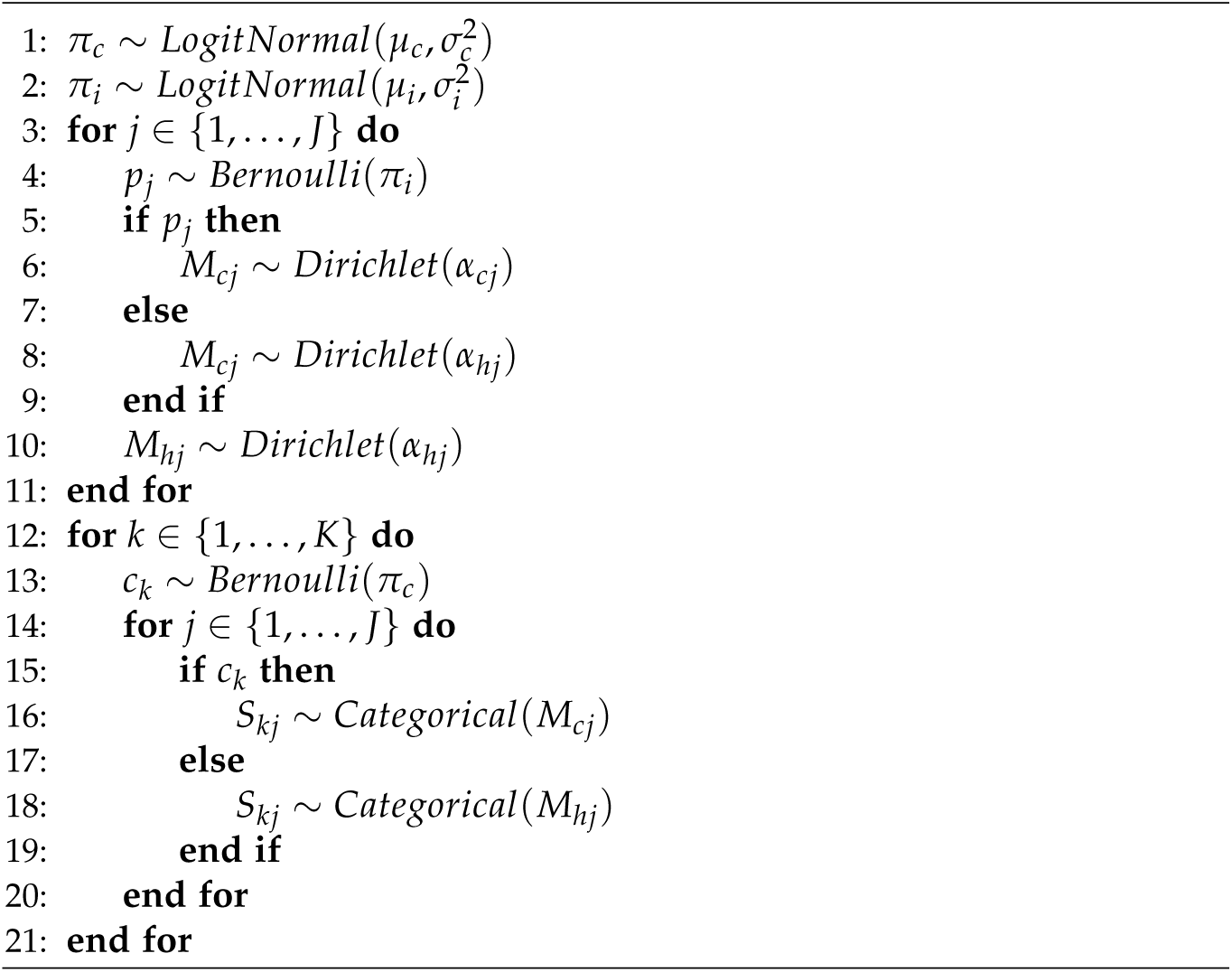

We performed inference using the generative model described by Algorithm 1 coupled with Metropolis-Hastings sampling on probabilistic program traces [25] using two gut microbiome databases, IGC [26] and our own assembly of the T1D DIABIMMUNE cohort [6]. Metagenes, microbial genes from a metagenome, were filtered to contain a valid open reading frame present in at least 10% of donors. The insulin chain INS B 9–25, a region the primary autoantigen in T1D has been mapped to [27], was used as a target epitope.

Tables 1 and 2 list the top 10 hits for each database assembly, sorted by posterior probability. The homology column uses the notation introduced by Clustal Ω [28], with a colon punctuation mark for highly similar but not identical amino acids, a dot for close substitutions and a space for unrelated amino acids. The BLOSUM45 matrix was used to define a distance metric among pairs of amino acids [29].

**Table 1:** Top predicted IGC bacterial mimotopes of INS B chain, sorted by posterior probability *p*.

| Metagene | Sequence | Homology | $p$ |
| --- | --- | --- | --- |
| 3043114 | GHSVEALYAVLAKEGFF | .H VEALY V .E:GFF | 0.93 |
| 3160029 | GHSVEALYAVLAKKGFF | .H VEALY V .:GFF | 0.90 |
| 3247770 | GHTVEALYAVLCQKGFF | .H VEALY V .:GFF | 0.89 |
| 3169624 | GHTVPALYAVLAKEGYF | .H V ALY V .E:G:F | 0.77 |
| 3161322 | GHSVEAYYAVLAKEGFF | .H VEA Y V .E:GFF | 0.75 |
| 2969430 | GHSVEAYYSVLAKEGFF | .H VEA Y V .E:GFF | 0.74 |
| 3278638 | GHSVEALYCILADRGFF | .H VEALY : .:RGFF | 0.73 |
| 3186877 | GHSVETLYCILADRGYF | .H VE:LY : .:RG:F | 0.72 |
| 3298615 | GHIAEALYVTLAKRGFF | .H: .EALY:. .:RGFF | 0.52 |
| 3205639 | GHIAEALYVVLAKKGYF | .H: .EALY:V .:G:F | 0.51 |
| INS B 9–25 | SHLVEALYLVCGERGFF | .H . : Y . .:G:F |  |

**Table 2:** Top predicted DIABIMMUNE bacterial mimotopes of INS B chain, sorted by posterior probability *p*.

| Metagene | Sequence | Homology | $p$ |
| --- | --- | --- | --- |
| 267926_71 | GHSVEALYAVLAKEGFF | .H VEALY V .E:GFF | 0.96 |
| 750513_07 | GHSVEALYAVLCDKGFF | .H VEALY V ::GFF | 0.88 |
| 778380_12 | GHTVEALYAVLCQKGFF | .H VEALY V ::GFF | 0.80 |
| 045379_02 | GHTVPALYAVLAKEGYF | .H V ALY V .E:G:F | 0.74 |
| 611716_02 | GHSVEAYYSVLAKEGFF | .H VEA Y V .E:GFF | 0.72 |
| 846083_03 | GHSVEALYCILADRGFF | .H VEALY : .:RGFF | 0.70 |
| 974697_44 | GHTVPALYATLAERGFF | .H V ALY . .ERGFF | 0.65 |
| 037280_07 | GHIAEALYVTLAKRGFF | .H: .EALY: .:RGFF | 0.56 |
| 947478_02 | GHIAEALYVVLAKKGYF | .H: .EALY:V .:G:F | 0.55 |
| 822586_01 | GHCAPALYAVLAERGFF | .H . ALY V .ERGFF | 0.54 |
| INS B 9-25 | SHLVEALYLVCGERGFF | .H . A Y . .:G:F |  |

The same pattern stands out in both databases, with 3 or perhaps 4 blocks of identity and homology, plus a few non-conserved regions. This motif degrades with decreasing posterior probabilities, which might imply imperfect mimotopes are biologically relevant, contributing with additive effects towards tolerance. It may also reflect an evolutionary process towards mimicry.

Tables 3 and 4 list mappings between IGC and DIABIMMUNE, but also to the UniProt protein database [30]. These have been constructed by finding metagene matches with at least 97% overall identity, no gaps and no substitutions within the mimicry region. The species column has been taken from the corresponding UniProt entry. Many sequences are common between both top predicted hits. Others have matching proteins in the alternative assembly which did not enter the corresponding top results due to using a 10% abundance cutoff. This might reflect taxa changes between infant and adult guts, geographical variation, or differences induced by sample collection and sequencing parameters.

**Table 3:**
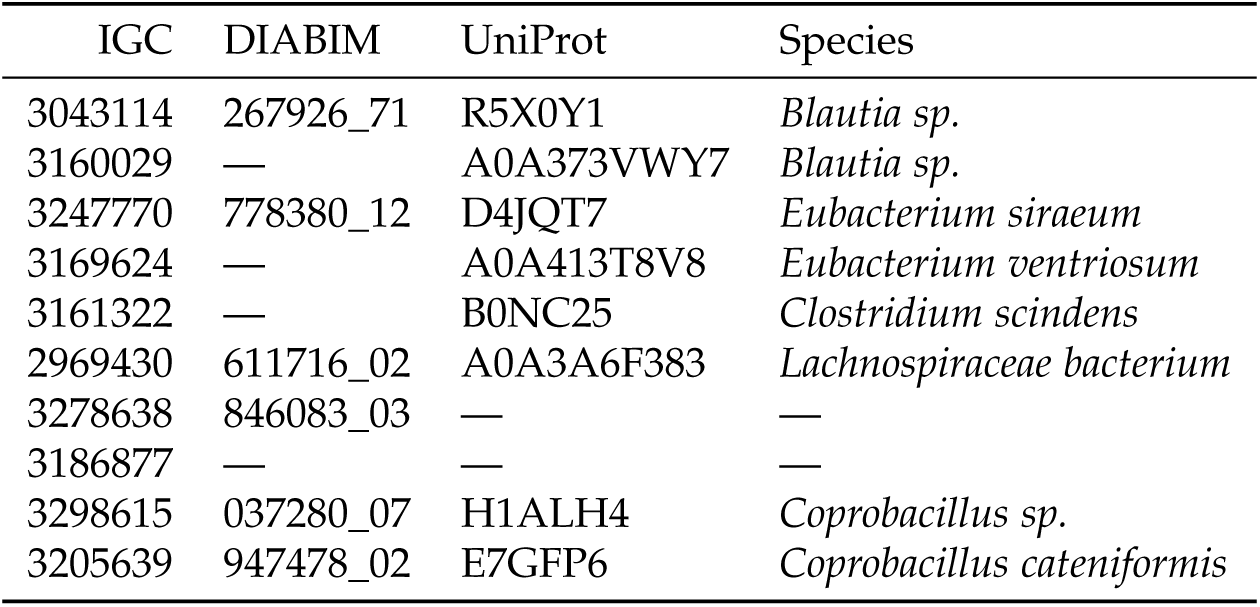
Mapping of top IGC bacterial mimotopes to DIABIMMUNE and UniProt.

| IGC | DIABIM | UniProt | Species |
| --- | --- | --- | --- |
| 3043114 | 267926_71 | R5X0Y1 | *Blautia sp.* |
| 3160029 | — | A0A373VWY7 | *Blautia sp.* |
| 3247770 | 778380_12 | D4JQT7 | *Eubacterium siraeum* |
| 3169624 | — | A0A413T8V8 | *Eubacterium ventriosum* |
| 3161322 | — | B0NC25 | *Clostridium scindens* |
| 2969430 | 611716_02 | A0A3A6F383 | *Lachnospiraceae bacterium* |
| 3278638 | 846083_03 | — | — |
| 3186877 | — | — | — |
| 3298615 | 037280_07 | H1ALH4 | *Coprobacillus sp.* |
| 3205639 | 947478_02 | E7GFP6 | *Coprobacillus cateniformis* |

**Table 4:** Mapping of top DIABIMMUNE bacterial mimotopes to IGC and UniProt.

| DIABIM | IGC | UniProt | Species |
| --- | --- | --- | --- |
| 267926_71 | 3043114 | R5X0Y1 | *Blautia sp.* |
| 750513_07 | — | — | — |
| 778380_12 | 3247770 | D4JQT7 | *Eubacterium siraeum* |
| 045379_02 | — | A0A413RAY3 | *Eubacterium ventriosum* |
| 611716_02 | 2969430 | A0A3A6F383 | *Lachnospiraceae bacterium* |
| 846083_03 | 3278638 | — | — |
| 974697_44 | 3268248 | A0A1M6LKK1 | *Megasphaera elsdenii* |
| 037280_07 | 3298615 | H1ALH4 | *Coprobacillus sp.* |
| 947478_02 | 3205639 | E7GFP6 | *Coprobacillus cateniformis* |
| 822586_01 | 2887265 | A0A174KQ89 | *Collinsella aerofaciens* |

Nonetheless, reproducibility is particularly noteworthy. All species listed belong to the Firmicutes phylum, except *Collinsella aerofaciens* which is classified as an Actinobacteria. Other Actinobacteria are present in entries that are located close to the highest posterior probabilities. By inspecting assembled proteins or the corresponding UniProt entry, listed hits appear to be highly variable in terms of sequence length and amino acid composition except within the mimotope region.

### 2.2 Insulin B mimotopes belong to the transketolase superfamily

In case of the IGC reference gut microbiome, 49 out of 50 hits with highest posterior probabilities are annotated as belonging to the Kyoto Encyclopedia of Genes and Genomes (KEGG) ortholog class K00615 [31]. This corresponds to the transketolase (TKT) enzyme domain, part of the pentose phosphate pathway. Given that K00615 is rare in IGC, with just 6176 entries out of a total 9879896 (0.06%), this result appears to be significant. A formal enrichment test confirms this fact with a p-value estimate < 10^−6^. Reannotating DIABIMMUNE yields equivalent results, given that 48 out of 50 entries containin domain K00615.

Pfam is a database that contains profile hidden Markov models (pHMMs) [32] derived from curated multiple alignments [33]. Within the Pfam pHMM for TKT, accession no. PF00456, all mimotope regions map to residues 63–79. Since insertions or deletions are very unlikely in that interval, we can treat the model as a simpler position-specific weight matrix (PWM). The latter describes the probability of each amino acid per position independently of other residues [34]. The lack of insertions, deletions and hidden states allows for a much easier mathematical treatment and visualization.

A sequence logo plot is derived by making the PWM plot y axis length at each residue equal to its information content [35]. Positions where all amino acids are equally likely will have a maximal entropy and carry no information at all. The logo plot in Figure 1 shows TKT residues in the region of interest are relatively well conserved across all superfamily members, whereas the others are highly random indicating they might not be required for the correct functioning of the enzyme. Amino acids occupying highly conserved residues and their spacing seem very well suited for evolution to develop mimotopes employing TKT as a template. This is also facilitated by the lack of conservation in the remaining residues, which can be mutated without disrupting the function of the enzyme. Given that insulin mimotopes encoded in TKT are present across two bacterial phylums, Firmicutes and Actinobacteria, horizontal or lateral gene transfer events are probable. Phylogenetic tree reconstruction supports the existence of these events [36], as well as some observations derived from insect models [37].

**Figure 1:**
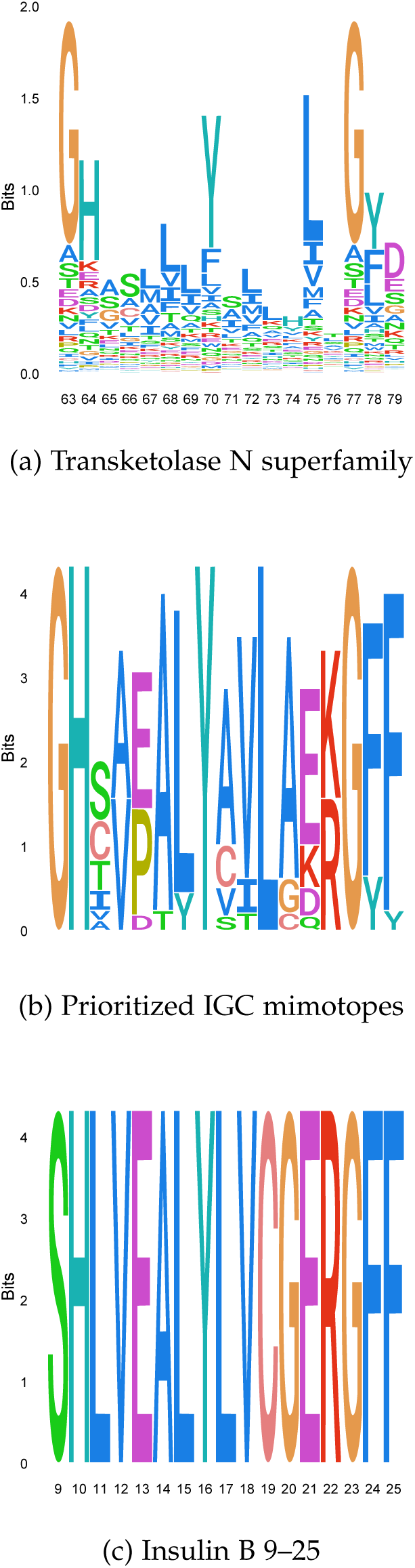
Sequence logo for transketolase N superfamily, prioritized IGC mimotopes and insulin B 9–25 chain.

### 2.3 Insulin B and transketolase mimotopes are crossreactive

To obtain further experimental evidence that supports these findings, we activated CD4 T cells using monocyte-derived dendritic cells as antigen presenting cells isolated from whole blood belonging to a diagnosed T1D donor. Synthesized peptides matching some of the top predicted IGC and DIABIMMUNE mimotopes, outlined in Table 5, were used. Four CD4 T cell pools were created. Pool U was not stimulated. Pool C was stimulated with a set of control peptides, pool I with all synthesized insulin peptides, and pool T with all mimotopes listed in Table 5.

**Table 5:**
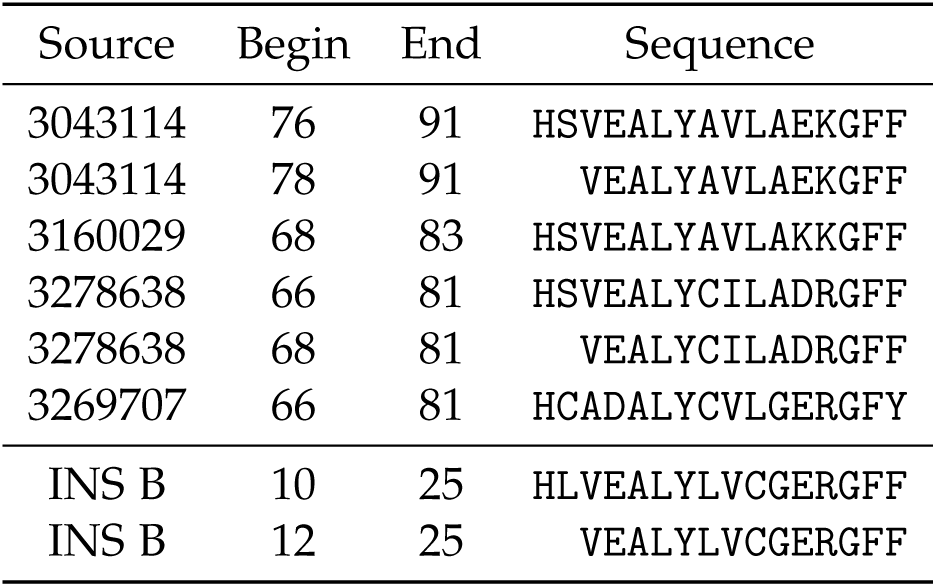
Mimotopes and insulin B chain peptides used to assess commensal crossreactivity.

**Table 6:**
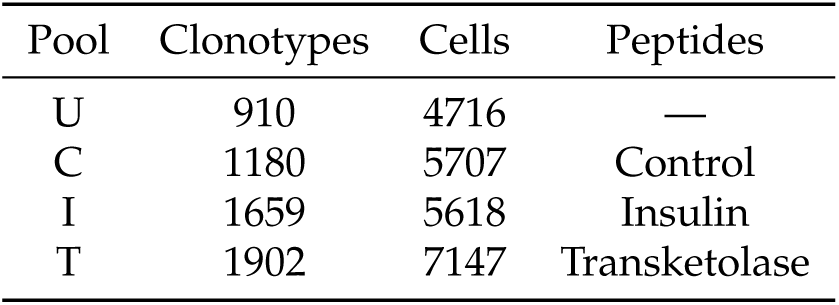
Monocyte-derived DC assay restimulation pools.

The intersection of clonotypes present in pools I and T with at least frequency 10, minus any clonotype present in any other pool at any frequency returned the 8 clonotypes listed in Table 7. All other equivalent pairwise intersections across pools returned an empty set or just one public clonotype. Not only a few high frequency clonotypes are shared and exclusive to the insulin and TKT pools, but some do have extremely low predicted recombination probabilities for β chains [38] and exhibit high degrees of sequence similarity suggesting equivalent binding properties.

**Table 7:**
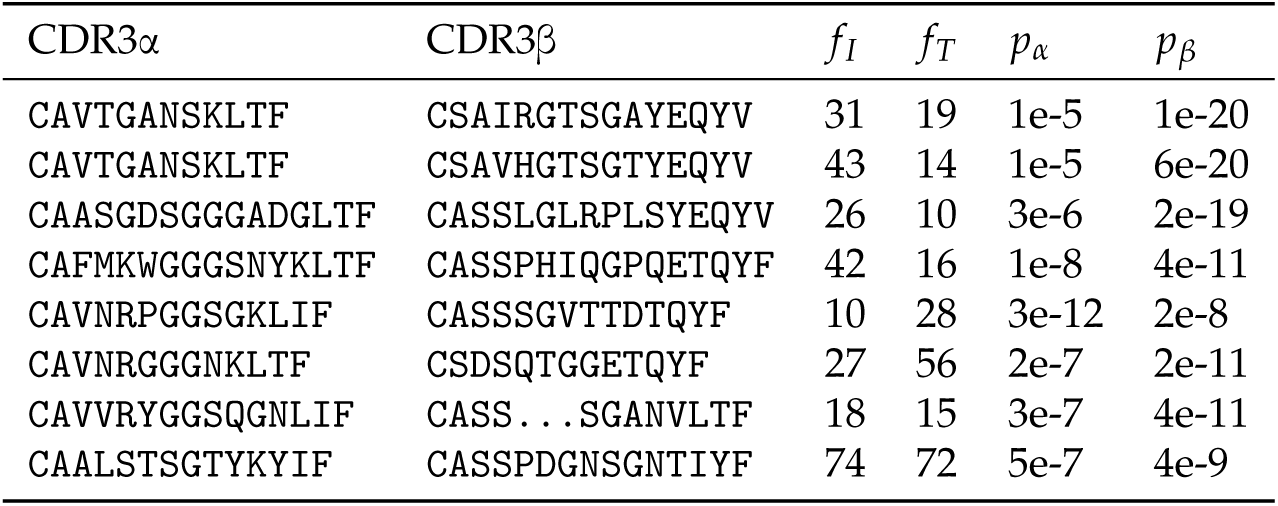
High frequency clonotypes stimulated by insulin B and mimotopes.

## 3 Discussion

The discovery that some bacterial TKT enzymes contain functional mimotopes of the key autoantigenic epitope in T1D, INS B 9–25, supports the hypothesis that self is partly encoded in the microbiome. In some scenarios, these peptides are probably mimicking self antigens recognized by a large number of regulatory T cells, as that is an optimal solution that confers bacterial communities maximal fitness. However, in most cases, mimotopes probably have less pronounced effects, as bacteria have not been able to adapt their sequences enough or target key self peptides. Hence, only the combined presentation of many weak mimotopes possibly recruiting few regulatory T cells each, to antigen presenting cells, would be enough for an overall tolerogenic self classification. This could also explain the difficulties in identifying mimotopes, as most sequences would exhibit weak mimicry motifs. Non-peptide mimotopes [39] may also have an underappreciated role in conveying signals currently denoted as immunological context [40].

Dysbiosis, defined as a disruption of commensal communities, alters the proportion of peptides and context signals presented. The mimotopes evolved by selection pressure are no longer sufficient to elicit a tolerogenic self response. This has important potential consequences for the host organism. Repeated presentations of the mimotope in a dysbiotic context could lead to an autoimmune response against the bacterial antigen. Which implies the self epitope it mimicks also becomes downgraded, as they are undistinguishable, and no longer has a positive contribution towards tolerance when presented along other self peptides. Changes of epitope status can be thought of, in practical terms, as for example alterations of T conventional versus regulatory cell proportions recognizing them.

Aside from molecular mimicry, there are several independent sources of evidence pointing to the role of TKT in T1D onset. The TEDDY study has shown TKT to be one of the most consistently upregulated microbial enzymes during the first year of life [23]. The TKT enzyme is involved in the non-oxidative branch of the pentose phosphate pathway and has been linked to the fermentation of fibre [41]. This becomes an important metabolic pathway after the transition to solid food, known as weaning, that typically occurs between 4 to 12 months of age. Microbial TKT has been also implicated in the metabolism of exopolysaccharides, high-molecular-weight polymers secreted by microbes into the microbial ecosystem, promoting a shift in the microbial metabolism to adapt to new dietary conditions and render more reducing power [42]. TKT expression confers microbes an important adaptation in sourcing energy, including saccharides produced by other microbes such as bifidobacteria, and thus allows rapid host colonisation [43].

Notwithstanding the benefits of fibre-rich diets are widely accepted, early introduction of fibre especially at time of dysbiosis may also propagate pre-existing TKT-expressing microbes and lead to T1D. Early introduction of root vegetables during infancy has been associated with increased risk of insulin autoimmunity in a cohort of children with high genetic susceptibility to T1D [44].

A recent study has demonstrated the importance of the intestinal microbiome inducing a vigorous immune response known as a weaning reaction [45]. While the weaning reaction is programmed in time to allow healthy immune development, at state of microbial dysbiosis the weaning reaction can lead to responses against mimotope-expressing commensal bacteria.

The TEDDY study has shown autoantibodies are rare before 6 months of life [46]. However, autoantibodies against insulin typically develop first shortly afterwards, with a highest incidence around 12 months of age, and a median to seroconversion of 18 months. Hence, both the time expression pattern of TKT and autoantibody development support the hypothesis of dysbiosis leading to immunity against the microbiome, and eventually against insulin due to mimicry. This process may be initiated before transition to solid food, and occur at a time when undeveloped epithelial barriers can cause microbial TKT leaks into the host [47].

In particular, dysbiosis might start before weaning if infants are fed artificial milk formulas lacking key human milk oligosaccharides that naturally support healthy metabolism, growth of bifidobacteria and gut homeostasis. TKT expression may be an advantage for host colonization as it allows specific microbes to efficiently source energy from polysaccharides present in the diet or from polysaccharides produced by other microbes. Therefore TKT may link microbial metabolism of polysaccharides, dysbiosis and antigen-specific immune responses to insulin.

This is also supported by the lower metagene abundance in T1D children after seroconversion and before diagnosis observed by DIABIMMUNE. Some of these changes seem to come from *Blautia, Rikenellaceae*, and the *Ruminococcus* and *Streptococcus* genera [6]. *Blautia* is the species containing the top insulin mimotope in both reference gut microbiome datasets, as shown in Tables 3 and 4. Gut dysbiosis has been associated with dysregulated abundance of essential short chain fatty acids, products of polysaccharide processing by microbes, and increased gut permeability or leaky gut syndrome [48]. Overall, our findings suggest that dysbiosis, linked to colonisation by TKT-expressing microbes, may destabilize epithelial barriers of the gut causing leak of mimotopes to the host followed by an autoimmune response to insulin.

Vitamin B1 (thiamine) deficiency is a frequent complication in T1D that leads to widespread tissue damage [49]. Thiamine is a well known co-factor of TKT [50], and its deficiency and low TKT activity are highly correlated. Notably, changes in gut microbiome taxa abundances and transcribed pathways of T1D patients have been shown to precede thiamine deficiency [51].

Probiotics and synbiotic supplementation improve peanut allergies [52] or reduce mortality in children with sepsis [53] by promoting gut microbial homeostasis. Our results impact future design and implementation of such strategies. It is possible that synbiotic supplementation would enhance tolerance induction to insulin in newborns previously stratified for genetic T1D susceptibility, where transketolase mimotopes would have to be taken into account. Finally, after decades of research, ever since particular HLA class II molecules were implicated as the strongest genetic risk factor [54], we present a possible missing major environmental link—dysbiosis-sensitive microbial mimotopes of insulin B 9–25 as a primary and specific cause of type 1 diabetes.

## Acknowledgements

This study is supported by a JDRF / Wellcome Strategic Award. Arcadio Rubio García acknowledges a PhD scholarship from JDRF and support from DARPA under its Probabilistic Programming for Advanced Machine Learning (PPAML) program. Dmech blood donors were recruited under the South Central Oxford A Ethics Committee, Ethics Reference 18/SC/0559, IRAS Project ID 243305. We would like to thank Katharine Owen and Rachel Besser, Oxford University Hospitals, for donor recruitment and members from the Diabetes and Inflammation Laboratory for blood processing: Heather McMurray, Shannah Donhou, Sarune Kacinskaite, Michael Ellis, Sandra Banks and Georgina Burton. We acknowledge Melanie Dunstan for additional help with sample processing; Olga Platonova, Claire Scudder, Raqeeb Mahmood and Sylwia Kopijasz for ethics management; and Dominik Trzupek, Jamie Inshaw, Hong Harper and Florent Yvon for genotype data queries. We would like to thank Moustafa Attar, Oxford Genomics Centre, for technical assistance with the 10X Chromium Single Cell genomics platform.

